# AMADEUS: Annotation-free multi-animal direction estimation using self-supervised learning

**DOI:** 10.64898/2026.09.19.752854

**Authors:** Yusuke Notomi, Kentarou Matsumura, Shigeto Dobata

**Author notes:** Correspondence: Y.N., S.D.

## Abstract

Markerless multi-animal tracking has advanced rapidly, yet measuring social interactions among visually similar animals remains challenging during occlusion and crowding. Here we present AMADEUS (Annotation-free Multi-Animal Direction Estimation Using Self-supervised learning), a tracking system that requires neither manual training annotation nor physical marking. AMADEUS extracts masks representing single animals, uses movement direction to assign head direction, and then synthesizes interaction images using copy-paste augmentation. A detector trained on the synthetic dataset estimates oriented bounding boxes and head direction, from which center, front and rear keypoints are derived. For a video containing instances of almost complete occlusion, AMADEUS additionally uses contrastive learning for identity verification. Across benchmark videos of 10 mice, 10 fish, 32 beetles, 32 ants with one cricket, 60 flies, and 368 ants, AMADEUS outperformed state-of-the-art methods in identity tracking and keypoint localization. AMADEUS enables scalable quantification of interactions across diverse species.

---

Quantitative analysis of animal behavior is fundamental to ecology, neuroscience, and ethology. Advances in video recording and computer vision have increased the scale and detail of behavioral measurements, creating growing demand for automated analysis 1,2. One important challenge in automated video analysis is measuring social interactions, which requires tracking multiple individuals while maintaining their identities over time. Individual identification often relies on color markings and physical tags 3–5, which allow direct and reliable identification of individuals as long as the markers remain detectable, but the markers can be obscured or lost and may alter behavior 5–7. Moreover, physical marking can become impractical when tracking large numbers of individuals or very small animals because of the effort required to mark each individual. These limitations of physical marking have motivated the development of markerless tracking methods.

A range of markerless approaches has been developed for multi-animal tracking. Many approaches, including FastTrack ^8^, idtracker.ai ^9,10^, TRex ^11^, UDMT^12^, and UMATracker ^13^, incorporate segmentation to localize animals. When animals cross or occlude one another, segmentation may fail to separate individuals, making identity tracking difficult ^10,12^. Object detectors based on deep learning, such as YOLO ^14^, can detect individual animals directly in images without segmentation, but conventional supervised training typically requires substantial manual training annotation.

Many multi-animal tracking systems primarily provide position and identity trajectories, but these outputs alone are not sufficient for many analyses of social behavior. DeepLabCut ^15^ and SLEAP ^16^ have enabled detailed pose estimation and analysis of interactions from video, but still require substantial manual training annotation for supervised training. Even essential pose information provided by front and rear keypoints greatly increases the information that can be obtained about social interactions, including approach direction and alignment. Existing annotation-free approaches such as UDMT ^12^, idtracker.ai ^9,10^ and UMATracker ^13^ focus primarily on position and identity, whereas FastTrack ^8^ and TRex ^11^ additionally derive essential pose information. However, because these estimates are derived from foreground components, the pose of each individual cannot be estimated when interacting individuals cannot be separated into distinct components. Reliable annotation-free identity tracking while retaining essential pose information during frequent crossings and crowding therefore remains challenging ^12^.

Here we present AMADEUS (Annotation-free Multi-Animal Direction Estimation Using Self-supervised learning), a markerless multi-animal tracking system for laboratory videos. AMADEUS extracts connected components of foreground pixels (blobs) and retains blobs each of which represent one complete and isolated individual as single-animal blobs. An oriented bounding box (OBB) is fitted to each single-animal blob. The class label associated with each OBB, which is ordinarily used to represent an object category, is instead used to represent head direction as one of eight direction classes. Building on copy-paste data augmentation for object detection and instance segmentation ^17–20^, AMADEUS synthesizes interaction images by copy-pasting single-animal blobs. A detector trained on the synthetic dataset estimates OBBs and direction classes, from which head directions expressed as continuous angles and center, front, and rear keypoints are derived. The user only needs to configure the foreground segmentation and answer the initial three questions: “Do the animals ever move backward?”, “How severe is the overlap?”, and “How many animals are in the video?” No manual training annotation or physical marking is required.

Across six benchmark videos with group sizes ranging from 10 to 368 animals, AMADEUS outperformed state-of-the-art methods in identity tracking and keypoint localization. To illustrate the behavioral analyses enabled by AMADEUS, we quantified pairwise alignment and the temporal sequence of turns in fish and mapped contact positions during interactions between host ants and an ant-loving cricket. The tracking system has been released as an open-source software package with a graphical user interface (GUI) designed for easy setup and operation. AMADEUS enables scalable multi-animal tracking across diverse species under conditions including frequent crossings and crowding.

## Results

### A self-supervised workflow generates synthetic interaction training data

AMADEUS provides an end-to-end graphical user interface for configuring and controlling the workflow, allowing each processing stage to be run, stopped, or skipped (Fig. 1). After loading a video, the user configures foreground segmentation using a real-time preview. Once the initial settings are specified, subsequent label generation, synthetic data construction, detector training, detection, and tracking proceed automatically as a batch process.

**Fig. 1.**
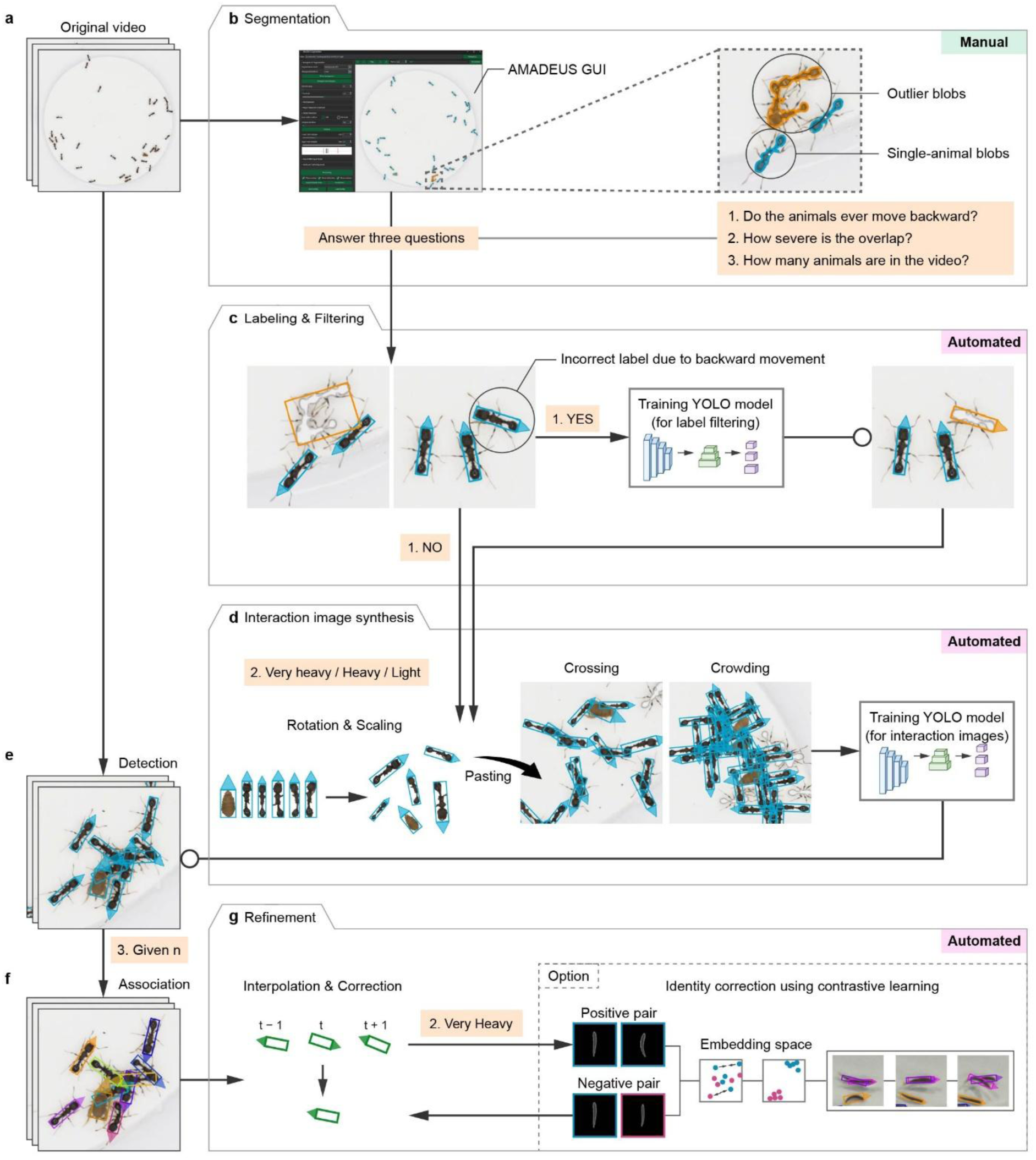
Overview of the AMADEUS workflow. **a**, Original video used as input, illustrated using the video of 32 ants with one cricket. **b**, Foreground segmentation after manual configuration. Cyan masks indicate single-animal blobs and orange masks indicate outlier blobs. **c**, Automatic labeling and filtering for single-animal blobs. An OBB is fitted to each single-animal blobs, and one of eight head direction classes is assigned from movement direction. When backward movement is expected, a YOLO OBB detector filters potentially incorrect class labels. The circled individual shows an incorrect label caused by backward movement. In c–g, triangles attached to the OBBs indicate head direction. **d**, Interaction image synthesis by copy-paste augmentation. Donor single-animal blobs are randomly rotated, scaled and pasted near target blobs to synthesize crossings and crowding. The synthetic images are used to train a YOLO OBB detector. **e**, Multi-animal detection using the detector trained on synthetic interaction images. The detector estimates OBBs and direction classes during crossings and crowding, including OBBs that approximate the overall body shape when only part of an animal is visible. **f**, Association of detected OBBs using Kalman filter prediction and Hungarian assignment. Colors denote individual identities. **g**, Refinement using geometric and temporal information. Short gaps are interpolated, and transient flips in head direction and positional inconsistencies are corrected. When almost complete occlusion is expected, contrastive learning is additionally used for identity correction. Images of isolated individuals aligned by head direction are used to learn an embedding space representing visual appearance, using positive pairs of images from the same individual and negative pairs from different individuals. The resulting embedding space is then used to detect and correct identity switches.

The workflow first extracts single-animal blobs for training (Fig. 1a, b). Blobs within the expected area range for single animals are retained, whereas blobs outside this range are classified as outlier blobs, including multi-animal blobs formed by merged individuals and fragmented blobs representing only part of an individual animal. For each remaining single-animal blob, an OBB is fitted, and one of eight classes representing head direction is assigned based on movement direction (Fig. 1c). The eight direction classes correspond to directions separated by 45°, such as “upper”, “upper left”, and “left”. Blobs are also treated as outliers when displacement between corresponding blobs of the same identity in consecutive frames is insufficient for reliable direction class assignment. Outlier blobs are replaced with background in the training images. When an animal moves backward, the assigned class can be incorrect. When backward movement is expected, a YOLO OBB detector is trained on images containing only the retained single-animal blobs to identify and exclude such potentially incorrect labels. Because the detector used to filter direction class labels is trained only on images containing single-animal blobs, the detector is used only to filter potentially incorrect labels and is not used in subsequent processing.

The images containing only single-animal blobs are then used as base images for copy-paste data augmentation (Fig. 1d). Donor single-animal blobs are randomly rotated and scaled and pasted near target blobs to synthesize interaction images involving two or three animals, as well as images of denser crowding configurations. Single-animal blobs to which no additional animals are pasted are also retained as training samples. Each video produces 10,000 images, comprising 9,500 training images and 500 validation images, which are used to train a YOLO OBB detector. The detector estimates OBBs and direction classes even with frequent crossings and crowding. Training on synthetic interaction images with partial occlusion enables the detector to estimate OBBs that approximate the overall body shape even when only part of an animal is visible (Fig. 1e).

Detected OBBs are then associated over time (Fig. 1f). Three Kalman filters predict OBB center, shape, and head direction, and Hungarian assignment ^21^ associates current detections with existing tracks using these predictions and the most recent valid OBBs. Geometric and temporal information is further used to recover short gaps and correct transient flips in head direction and positional inconsistencies (Fig. 1g). When the user indicates during the initial question that almost complete occlusion is expected, contrastive learning is additionally used. Identity switches are detected and corrected by comparing images of isolated individuals aligned by head direction before and after interactions. The final output provides individual identity, OBB center and shape, head direction, and derived front and rear keypoints over time.

### Benchmark videos and methods used for comparison

We evaluated AMADEUS on six benchmark videos of 10 mice, 10 fish, 32 beetles, 32 ants with one cricket, 60 flies, and 368 ants (Fig. 2a). In the videos of 10 mice and 10 fish, frequent crossings and substantial body deformation occurred. In the video of 32 beetles, individuals were relatively well separated, but periods of little or no movement and instances of backward movement limited the reliability of movement direction for class assignment. The video of 32 ants with one cricket included frequent crossings and two morphologically distinct species. DeepLabCut and SLEAP required separate training datasets and models for the ants and the cricket, whereas AMADEUS analyzed both with a single detector. Dense aggregations in the videos of 60 flies and 368 ants challenged foreground separation, detection, identity tracking, and computational scalability.

**Fig. 2.**
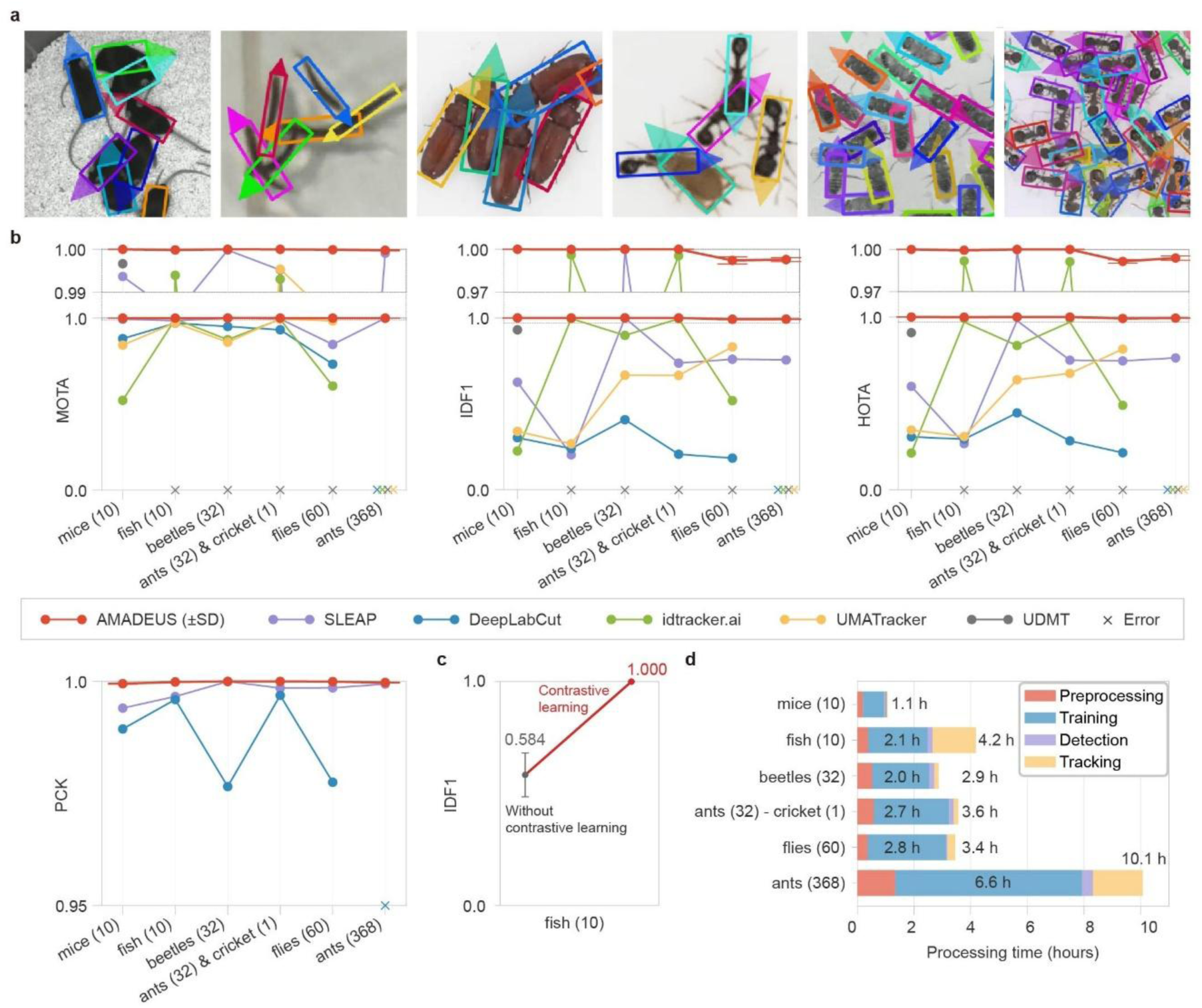
Benchmark comparison of AMADEUS. **a**, Representative AMADEUS tracking results for six benchmark videos. From left to right, the benchmark videos contained 10 mice, 10 fish, 32 beetles, 32 ants with one cricket, 60 flies, and 368 ants. Triangles attached to the OBBs indicate the head direction. **b**, Comparison of the final AMADEUS output after refinement (red) with outputs from SLEAP (purple), DeepLabCut (blue), idtracker.ai (green), UMATracker (yellow), and UDMT (gray). Post-processing procedures provided by each method were applied where available. Multiple Object Tracking Accuracy (MOTA), ID F1 score (IDF1), Higher Order Tracking Accuracy (HOTA), and Percentage of Correct Keypoints (PCK) are shown. Expanded views near 1.0 are shown above the MOTA and IDF1 plots. **c**, Effect of identity correction using contrastive learning on IDF1 in the video of 10 fish. Identity correction was applied only to the fish benchmark. **d**, Processing time for preprocessing (red), training (blue), detection (purple), and tracking (yellow) across the six benchmark videos. For the video of 10 fish, the time for tracking includes the time required for identity correction using contrastive learning. Numbers at the right indicate the total processing time.

We compared AMADEUS with DeepLabCut, SLEAP, idtracker.ai, UDMT, and UMATracker. DeepLabCut and SLEAP were trained separately for each benchmark video using manually annotated front, rear, and additional keypoints selected for each species. Additional keypoints were included because using more keypoints than required for analysis can improve pose estimation performance ^15^. Approximately 8,000 keypoints were annotated for the video of 10 mice, 6,000 for the video of 10 fish, 12,800 for the video of 32 beetles, 13,200 for the video of 32 ants with one cricket, 24,000 for the video of 60 flies, and 14,720 for the video of 368 ants, totaling approximately 78,720 manually annotated keypoints. AMADEUS used none of these annotations and instead automatically generated OBB and class labels after manual configuration of the initial foreground segmentation. UDMT completed tracking only for the mouse video and terminated with errors for the other five videos. For the video of 368 ants, SLEAP was the only method other than AMADEUS that produced valid output. The other methods either terminated with errors or produced incomplete tracking results.

Ground truth (GT) was generated independently of AMADEUS output by manually correcting DeepLabCut output for the video of 32 ants with one cricket and SLEAP output for the other videos. Manual correction removed clear tracking and keypoint errors, but small residual deviations in front and rear keypoints were not exhaustively corrected. To reduce sensitivity to these residual GT errors and to avoid favoring or penalizing the methods whose outputs were used to initialize the GT, a tolerance of 0.5 body length was used for evaluation.

### AMADEUS maintains accurate positions and identities

We evaluated position and identity tracking using Multiple Object Tracking Accuracy (MOTA) ^22^, Higher Order Tracking Accuracy (HOTA) ^23^, and ID F1 score (IDF1) ^24^. For these metrics, the midpoint between the front and rear keypoints was used as the position for AMADEUS, SLEAP, and DeepLabCut. For idtracker.ai, UMATracker, and UDMT, the reported position was used instead. A predicted position was matched to a GT position when the distance between them was within 0.5 body length. To use the same matching threshold across metrics, HOTA was calculated at a single threshold of 0.5 body length rather than averaged across multiple thresholds ^23^. MOTA, HOTA, and IDF1 therefore measure different aspects of tracking performance using the same matching threshold.

Performance was evaluated using the final outputs from each method, including the post-processing provided by each software package where applicable. For the fish video, identity correction using contrastive learning was additionally used because almost complete occlusion was expected. Across three random seeds, AMADEUS achieved mean MOTA values from 99.976% to 100.000%, HOTA values from 99.149% to 100.000%, and IDF1 values from 99.218% to 100.000% across the six benchmark videos (Fig. 2b). For every benchmark, AMADEUS outperformed every other method that produced valid output, with higher scores in all three metrics.

The difference between methods was particularly clear in identity tracking metrics under dense conditions. For the video of 60 flies, AMADEUS achieved 99.149% HOTA and 99.218% IDF1, whereas UMATracker, the strongest among the others for these metrics in this video, achieved 83.161% HOTA and 83.119% IDF1. For the video of 368 ants, MOTA was close to the maximum for both AMADEUS and SLEAP, at 99.976% and 99.909%, respectively. In contrast, AMADEUS achieved 99.391% HOTA and 99.279% IDF1, compared with 77.974% HOTA and 75.535% IDF1 for SLEAP. Thus, under the condition with the highest density, the larger performance difference was in maintaining individual identity rather than in the overall MOTA score.

AMADEUS also achieved high tracking accuracy under the other benchmark conditions. AMADEUS reached 100.000% MOTA, HOTA, and IDF1 for the beetle video despite instances in which movement direction did not reliably indicate head direction. For the video of 32 ants with one cricket, mean MOTA, HOTA, and IDF1 all exceeded 99.998%, despite frequent interactions between the two species. In the video of 10 mice, all three metrics exceeded 99.998%. These results demonstrate that AMADEUS can maintain accurate localization and identity despite substantial variation in body shape, movement, density, and species composition.

### Identity correction using contrastive learning resolves identity switches

The fish video revealed a limitation of identity tracking based on spatial, geometric, and temporal information. Although MOTA remained high, indicating that detections were largely complete and accurate (Fig. 2b), identity switches caused by almost complete occlusion persisted after the overlapping animals moved apart (Extended Data Fig. 1). These persistent identity switches could not be resolved by the standard tracking procedure based on spatial, geometric, and temporal information (Fig. 2c).

We therefore applied identity correction using contrastive learning to the video of 10 fish. The identity correction procedure was adapted from idtracker.ai v6 ^10^ and is implemented in AMADEUS. Images of individuals not in contact with other animals were aligned by head direction and used to learn an embedding space. The resulting embedding space was then used to verify identity assignments when association could not reliably maintain identity, correcting persistent identity switches (Fig. 2c). In the final output for the fish video, AMADEUS achieved 99.984% MOTA, 99.940% HOTA, and 99.986% IDF1.

These results indicate that association can maintain identity as long as sufficient spatial, geometric, and temporal information remains available, but persistent identity switches can arise when this information becomes insufficient during interactions. In AMADEUS, identity correction using contrastive learning is enabled when the user indicates during the initial setup that almost complete occlusion is expected. In such cases, identity verification using contrastive learning incorporates appearance information as an additional cue to restore identity continuity.

### Front and rear keypoint estimation

We next evaluated the front and rear keypoints derived from the OBBs and head directions estimated by AMADEUS. Head direction was expressed as a continuous angle. For each estimated OBB, the midpoint of the short edge in the estimated head direction was defined as the front keypoint, and the midpoint of the opposite short edge was defined as the rear keypoint. Localization accuracy was evaluated using Percentage of Correct Keypoints (PCK). Head direction was evaluated using the number of front–rear flips (FLIP). A FLIP was counted when an estimated position was matched to a GT position and the angular difference between the estimated and GT directions from the rear keypoint to the front keypoint was at least 90°.

Across the six benchmark videos, mean PCK ranged from 99.950% to 100.000% (Fig. 2b). Among videos with available DeepLabCut or SLEAP results, AMADEUS achieved higher PCK and lower FLIP counts in every comparison (Extended Data Fig. 1). For example, in the video of 368 ants, PCK remained high for both AMADEUS and SLEAP, at 99.976% and 99.942%, respectively, but the mean FLIP count was 485 for AMADEUS compared with 1,796 for SLEAP.

FLIP counts in AMADEUS were concentrated in the videos of 10 mice and 368 ants. In the video of 10 mice, strong body curvature can make the OBB nearly square, reducing the stability of the orientation of its long axis. In the video of 368 ants, extreme crowding occasionally degraded detection and head direction estimation. Nevertheless, across the diverse animal shapes and densities tested, AMADEUS maintained consistently high PCK and lower FLIP counts than DeepLabCut and SLEAP, without requiring manual keypoint annotation (Fig. 2b, Extended Data Fig. 1).

### Computational performance across benchmark videos

We next examined how processing time for AMADEUS was distributed across the workflow on a workstation equipped with an Intel Core i5-12400F CPU, 64 GB of RAM, and an NVIDIA GeForce RTX 4070 GPU with 12 GB of VRAM (Fig. 2d). The benchmark videos differed substantially in both group size and frame size (0.4–5.1 megapixels [MP]). Complete analysis required 1.1 to 10.1 h across the six benchmark videos. Detector training required 0.7 to 6.6 h and was the largest single component in all six videos. Detection speed ranged from 7.3 to 97.2 frames s⁻¹ and was generally lower for videos with larger frame sizes. For example, detection was approximately sixfold faster in the video of 10 mice (0.4 MP) than in the videos of 10 fish, 32 beetles, and 32 ants with one cricket (4.7–5.1 MP), while the lowest speed was observed in the video of 368 ants (4.7 MP).

The processing speed of tracking was approximately 52 frames s⁻¹ for the video of 10 mice (0.4 MP), 2 frames s⁻¹ for the video of 10 fish (5.1 MP), approximately 18 frames s⁻¹ for the videos of 32 beetles and 32 ants with one cricket (4.7 MP each), 10 frames s⁻¹ for the video of 60 flies (0.6 MP), and 2 frames s⁻¹ for the video of 368 ants (4.7 MP). Thus, except for the fish video, the processing speed of tracking generally decreased as the number of individuals increased, from 52 frames s⁻¹ for 10 mice to 2 frames s⁻¹ for 368 ants. For the fish video, tracking included identity correction using contrastive learning and required 1.5 h, accounting for 36% of the total processing time. Because identity correction using contrastive learning is intended for videos in which almost complete occlusion is expected, identity correction can be omitted when such occlusion is unlikely.

For comparison, the video of 10 mice was also processed on a laptop equipped with an Intel Core i7-1165G7 CPU, 32 GB of RAM, and an NVIDIA GeForce GTX 1650 Ti GPU with 4 GB of VRAM, requiring 4.8 h in total. Total processing time increased from 1.1 h on the workstation to 4.8 h on the laptop, largely because detector training increased from 0.7 to 4.0 h.

### AMADEUS enables analysis of collective behavior and social interactions

AMADEUS provides individual identity, OBB center and shape, head direction, and derived front and rear keypoints, enabling collective behavior and social interactions to be analyzed without additional annotation. We used AMADEUS to quantify pairwise alignment and sequential turns in the video of 10 fish, compare head directions inferred from centroid trajectories with head directions estimated by AMADEUS in the video of 32 beetles, and map contact positions during interactions in the video of 32 ants with one cricket (Fig. 3).

**Fig. 3.**
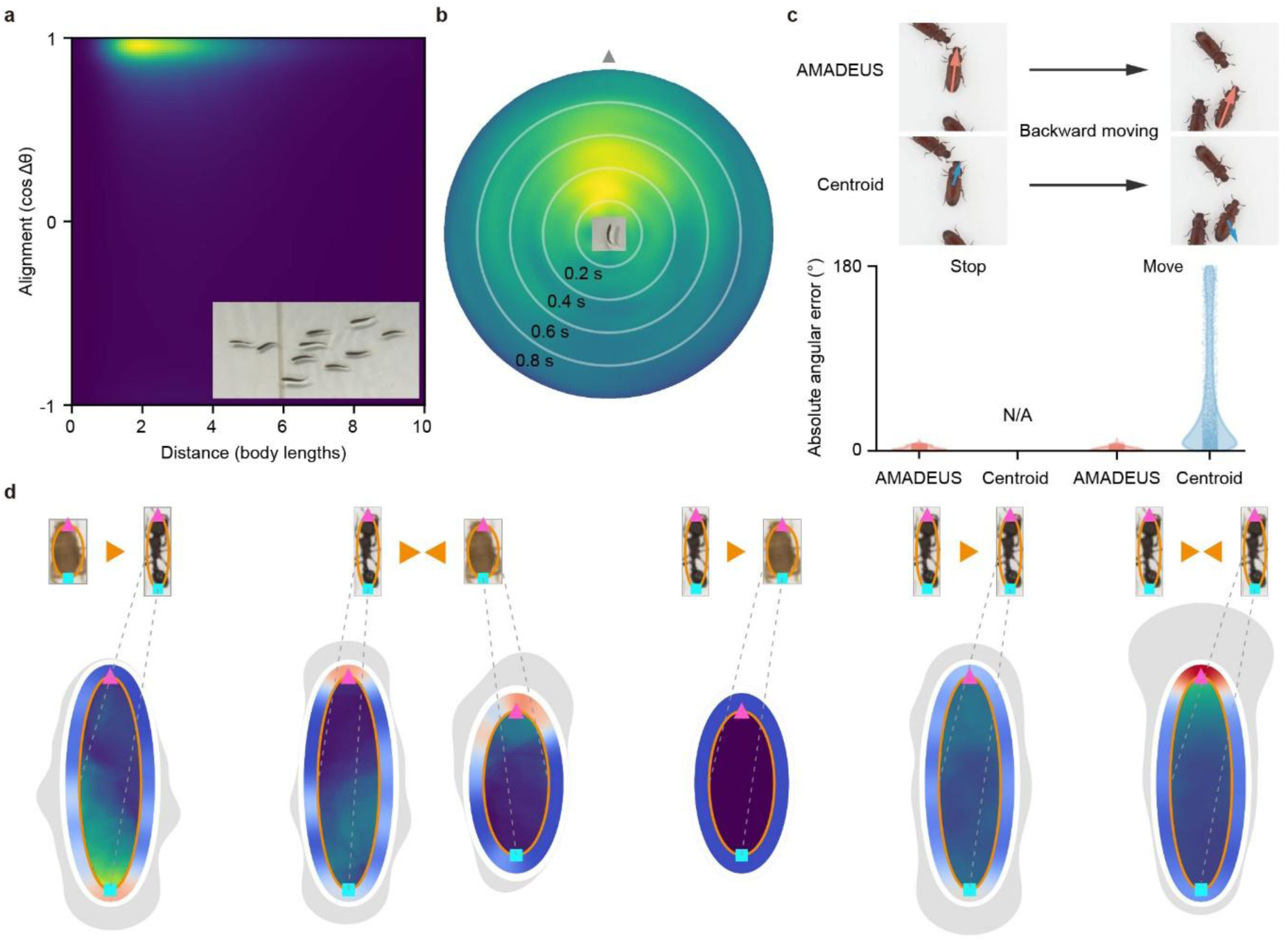
Behavioral analyses using AMADEUS output. **a,b**, Analyses of the video of 10 fish. a, Joint density of pairwise distance and alignment. Distance is normalized by the average of the mean OBB long side lengths of the two individuals. Alignment is the cosine of the difference between their head directions, with 1 indicating parallel and −1 indicating opposite directions. The image at the lower right shows a representative fish aligned with its head direction. **b**, Distribution of fish that turned earlier relative to a focal fish that subsequently turned, shown by response lag (radius) and relative direction from the focal fish (angle). Concentric circles indicate 0.2 s intervals, and the gray triangle marks 0°. **c**, Comparison of head direction estimated by AMADEUS and direction inferred from displacement of the GT centroid in the video of 32 beetles. Top, example of a beetle moving backward. AMADEUS estimates head direction during backward movement, whereas centroid displacement indicates movement direction. Bottom, absolute angular error relative to the GT head direction for stopping and moving animals. Head direction from centroid displacement is N/A while animals are stopping. **d**, Contact positions during interactions between ants and a cricket, classified by approach direction. A single orange triangle indicates a unidirectional approach and paired triangles indicate a mutual approach. Contact maps are shown relative to the focal animal using ellipses derived from detected OBBs. The orange outline indicates the mean body ellipse, and the magenta triangle and cyan square indicate the front and rear keypoints. The inner heat map shows mean overlap (or underlap) duration per contact. The colored ring and gray outer profile show circular kernel density estimates of contact onset positions, with equal weight for each contact and with weight proportional to contact duration, respectively.

In the fish video, we examined spatial alignment and the temporal sequence of turns among neighboring individuals (Fig. 3a,b). Pairwise distance was normalized by the arithmetic mean of the body lengths of the two individuals, and alignment was defined as the cosine of the difference between the head directions of the two individuals. The joint density of pairwise distance and alignment was concentrated at separations of approximately one to three body lengths and alignment values near 1, indicating predominantly parallel head directions (Fig. 3a). We then examined sequential turns in the same direction among neighboring fish. Turns by individuals ahead tended to be followed by turns in the same direction by individuals behind, with subsequent turns concentrated at lags of approximately 0.1 to 0.6 s (Fig. 3b). These analyses demonstrate that AMADEUS can characterize the spatial alignment and the temporal sequence of turns among neighboring fish even during frequent interactions.

In the beetle video, we then compared head directions estimated by AMADEUS with directions inferred from displacement of the GT centroid (Fig. 3c). AMADEUS showed low angular error for both stopping and moving animals, whereas centroid displacement could not provide head direction estimates while animals were stopping and showed substantial angular errors while animals were moving. For moving animals, the median angular error for each individual was significantly lower with AMADEUS than with centroid displacement (paired Wilcoxon signed-rank test, *n* = 32, *P* = 4.66 × 10^−10^).

Mapping contact positions relative to the head direction of each animal revealed differences associated with approach direction (Fig. 3d). During unidirectional approaches by the cricket, contact onsets were concentrated near the rear of the ant, with additional contacts along the sides of the ant, and mean overlap (or underlap) duration was also concentrated near the rear. During mutual approaches between an ant and the cricket, contact onsets were concentrated toward the front of both animals. Among contacts between ants, unidirectional approaches produced a broader distribution of contact onsets with a peak near the rear, whereas mutual approaches were concentrated toward the front. Combining OBB shape with head direction enabled contact positions to be mapped relative to the front, rear, and sides of each animal.

## Discussion

AMADEUS combines identity tracking with head direction estimation of animals in videos with frequent crossings and crowding, without manual training annotation or physical marking. The advance lies in interaction image synthesis from automatically labeled single-animal blobs, enabling detector training for conditions in which segmentation cannot reliably separate individuals. Across the six benchmark videos, AMADEUS achieved higher identity tracking accuracy than the other methods evaluated and more accurate front and rear keypoint estimates than DeepLabCut and SLEAP.

AMADEUS estimates an OBB and head direction for each animal, providing individual identity, OBB center and shape, head direction, and derived front and rear keypoints after tracking. The OBBs and head directions support analyses beyond position trajectories, as illustrated by pairwise alignment and sequential turns in fish and by contact positions in ants and a cricket. The behavioral analyses demonstrate how interactions can be quantified relative to the body of each animal. Moreover, front and rear keypoints are derived geometrically from OBBs, and contact is defined by overlap between OBBs or ellipses derived from OBBs. Although additional direct confirmation is needed to establish precise anatomical landmarks or physical contact, the essential pose information provided by AMADEUS can support a range of analyses of social interactions.

The following considerations should be kept in mind when applying AMADEUS. First, automatic label generation requires enough complete single-animal blobs for training and sufficient movement that indicates head direction. Persistent overlap or movement that rarely corresponds to head direction can limit automatic label generation. Strong body curvature can also destabilize the OBB long axis, as observed in the mouse video. Likewise, recording conditions should be designed to provide sufficient image quality for reliable foreground separation and head direction assignment based on movement. Nevertheless, compared with conventional tracking software based on foreground extraction ^8,10–13^, AMADEUS provides more options for foreground segmentation to accommodate moderate shadows and background fluctuations as seen in the video of 10 fish, as well as backgrounds containing regions with different levels of contrast against the animals as seen in the video of 10 mice.

AMADEUS is provided with a graphical user interface, with automatic processing once initial settings and foreground segmentation are configured. Although evaluated here only on animals, AMADEUS could also in principle be applied to motile cells or other rod-shaped moving objects. Application to such objects would require reliable foreground separation of isolated objects and movement that provides a reliable indication of head direction. When both requirements are met, AMADEUS may facilitate quantitative analysis across a broader range of species, experimental conditions, and videos without manual training annotation. Robust tracking during frequent crossings and crowding is particularly useful for studying dense groups, where frequent social interactions complicate tracking. By combining annotation-free training with robust identity tracking and spatial information about body orientation and shape, AMADEUS enables quantification of a wider range of moving objects and interactions from videos recorded in laboratory environments

## Methods

### Overview of AMADEUS

AMADEUS estimates oriented bounding box (OBB) and head direction of each animal in videos of visually similar, unmarked animals that frequently come into contact and cross paths, without requiring manually annotated training data. The workflow comprises (1) foreground segmentation; (2) association of single animal blobs; (3) selection of blobs for direction class assignment; (4) filtering of potentially incorrect class labels; (5) interaction image synthesis; (6) construction of training dataset; (7) training and detection; (8) multi-stage assignment; (9) refinement; and (10) optional identity correction using contrastive learning. Before processing, users specify the number of animals, the expected level of overlap, and whether backward movement occurs.

### Foreground segmentation

Foreground segmentation was the only processing stage requiring manual parameter adjustment. The segmentation threshold was adjusted using a real-time preview until the main bodies of isolated animals were extracted (Fig. 1b). A static background was estimated from the temporal median, maximum, or minimum at each pixel. Five segmentation modes were supported: background difference, dark-region thresholding, bright-region thresholding, and background difference intersected with either a dark-region or bright-region mask. The segmentation mode and region of interest (ROI) used for each benchmark video are listed in Extended Data Table 1.

Blobs were extracted with OpenCV 4.10.0.84 and classified as single-animal or outlier blobs. Outliers were identified using manually configured lower and upper thresholds for blob area (Extended Data Table 1). They included large blobs formed by animals in contact or overlapping, and small or fragmented blobs caused by debris, reflections, or incomplete segmentation. Blobs below a user-defined minimum area (Extended Data Table 1) were removed from the foreground mask as noise.

### Association of single-animal blobs

Single-animal blobs were associated across frames to generate tracklets, each comprising a sequence of associated blobs used to assign head direction based on movement. Blobs formed by animals in contact were classified as outliers and excluded from this association. The blob centroid and minimum-area OBB were computed for each single-animal blob. Mean body length was estimated from OBB long-side lengths after excluding values outside the Tukey bounds Q1 − 1.5 IQR and Q3 + 1.5 IQR, where Q1 and Q3 were the first and third quartiles of these lengths, respectively, and IQR was the interquartile range, defined as Q3 − Q1. Centroid displacement was expressed in units of this mean body length.

To select tracking parameters automatically, up to 100 windows of three consecutive frames were sampled evenly throughout the video. Within each window, single-animal blobs in consecutive frames were matched by Hungarian assignment to minimize the distance between blob centroids. Candidate pairs required a centroid distance of at most 1.5 mean body lengths and a ratio of later to earlier OBB long-side length between 0.5 and 2.0. These matches were used only for parameter selection, not to assign track identifiers (IDs). OBB intersection over union (IoU) was calculated as the intersection area divided by the union area. The association IoU threshold was the highest of 0.3, 0.4, and 0.5 met by at least 95% of matched pairs, or 0.3 if none qualified.

Single-animal blobs in consecutive frames were then associated using this IoU threshold. Candidate pairs meeting the threshold were accepted in descending order of IoU, with each blob and tracklet assigned at most once. Unmatched blobs started new tracklets. The same threshold was subsequently used to associate OBBs detected by the YOLO OBB detector trained on synthetic interaction images.

### Selection of blobs for direction class assignment

A blob was excluded from labeling if it was not associated with a tracklet, its OBB aspect ratio was below 1.1, or distance between its blob centroid and that of the preceding blob in the same tracklet exceeded twice the mean body length. OBB aspect ratio was defined as long-side length divided by short-side length. Each sequence of consecutive frames within a tracklet was also required to exceed a minimum duration. This duration was determined from the video frame rate and the 90th percentile of normalized centroid displacements in the matches used for automatic parameter selection. It corresponded to the time required to move one mean body length at this rate, rounded to 0.01 s and constrained to 0.20–0.50 s. Excluded and outlier blobs were replaced with background. Images containing only retained single-animal blobs, each with an OBB label, were generated every five frames.

Head direction was inferred from movement direction. Within each sequence, consecutive OBB long-axis vectors were first oriented consistently over time. A sequence was excluded if the distance between its first and last blob centroids was less than one mean body length. Local movement vectors were calculated from blob-centroid positions. For each of the two possible long-axis orientations, movement distance was weighted by the cosine of the angle between the long axis and local movement direction, then summed over the sequence. The orientation with the larger sum was assigned as the head direction and classified into one of eight direction classes separated by 45°. Because this procedure assumes that movement direction indicates head direction, potentially incorrect labels caused by backward movement were filtered in the next stage.

### Filtering of potentially incorrect class labels

When the user indicated that the video contained backward movement, a YOLO OBB detector was used to identify potentially incorrect direction class labels (Fig. 1c, Extended Data Table 2). All images containing only single-animal blobs were used to construct the training and validation dataset, with each OBB assigned one of the eight direction classes as its object class. The trained detector was applied to all images at a confidence threshold of 0.2. Each labeled OBB was matched to the overlapping detected OBB with the greatest IoU. A label was flagged if no match was found, the greatest IoU was below 0.80, or the detected and assigned direction classes differed by at least two 45° steps. Class differences were measured cyclically, so classes 1 and 8 differed by one step.

Within each tracklet, a sequence of consecutive frames was excluded if at least 40% of its blobs were flagged. Otherwise, the entire sequence was retained, including flagged blobs. Blobs in excluded sequences were replaced with background.

### Interaction image synthesis

AMADEUS generated synthetic interaction images for self-supervised learning by copy-pasting labeled single-animal blobs (Fig. 1d). Each image containing only single-animal blobs served as a base image. Donor blobs to be pasted onto base images were extracted using the segmentation masks. The blobs were allocated to configurations of one, two, or three animals in proportions of 0.1, 0.5, and 0.4, respectively. For two- and three-animal interactions, one or two donor blobs, respectively, were pasted near each target blob.

Donor geometry and appearance were randomly augmented before pasting. Rotation angles were sampled uniformly from 0° to 360°, and scale factors uniformly from 0.9 to 1.1. Additional, independent scaling by factors of 0.9–1.1 was applied along the OBB long axis only. Brightness and contrast were each adjusted by factors of 0.90–1.10. Segmentation masks used for pasting were expanded by factors sampled uniformly from 1.0 to 1.05. Donors were excluded if the OBB aspect ratio derived from the transformed mask or the visible mask after occlusion was below 1.1. The overlap area with the target had to be at least 0.01 times the donor mask area and at most 0.2 or 0.5 times that area, depending on the configured overlap level (Extended Data Table 2). Overlap with any other blob could not exceed the same maximum. Up to 100 placement attempts were made per donor to satisfy all conditions. The donor was then pasted either without masking, placing it in front of the target, or with the target-overlapping region masked out, placing it behind the target. These occlusion orders were equally probable.

To increase interaction diversity, additional groups of two or three donor blobs were generated independently of the original targets. For videos in which animals occupied a spatially localized region, as defined below, these groups were pasted into available empty space, with the number of groups approximately equal to the configured number of animals. For other videos, the number of additional groups was approximately 25% of the configured number of animals.

Clustered pasting generated dense configurations of up to 12 animals, including the target; failed donor placements resulted in smaller clusters. Because each original blob in a source frame could serve as a target, a synthetic image could contain multiple clusters and more than 12 animals in total. Donor OBBs were placed adjacent to OBBs already in the cluster. For placement calculations only, both dimensions of the target and donor OBBs were scaled to 80%, and donors were positioned so that the scaled OBB edges touched. Donor head directions were distributed as evenly as possible among parallel, antiparallel, and the two perpendicular directions relative to the target. Because conversion to pixel masks could produce slight overlap even between touching edges, the maximum permitted overlap was the greater of 8 pixels and 3.5% of the scaled donor OBB area.

### Construction of training datasets

The same image preparation procedure was used for the label-filtering detector and the detector trained on synthetic interaction images. Each source image yielded resized full-frame samples, high-resolution square crops, or both. The training image size was 1,024 pixels when the shorter frame dimension was at least 1,200 pixels; otherwise, it was the shorter dimension rounded down to a multiple of 32. Full-frame samples were resized with their aspect ratio preserved and padded to a square using the background estimated during segmentation. Crop number depended on the spatial distribution of isolated animals. If the union of original single-animal blobs fitted within one training-size square in at least 90% of valid frames, one crop centered on that union was generated. Otherwise, the numbers of training-size squares fitting along the frame width and height were multiplied, and the result was capped at four. Counts of one or two yielded one random crop; counts of three or four yielded two.

A crop label was retained only if at least 95% of the blob’s OBB area remained inside the crop. Otherwise, the region defined by the blob’s segmentation mask was replaced with estimated background, except where it overlapped the mask of a retained blob. This exception prevented erasure of retained animals. The replacement boundary was blended with the original image using Gaussian blending with a 7-pixel kernel and σ = 11.

For the label-filtering detector, 5% of samples were reserved for validation, with no fixed total sample count. For the detector trained on synthetic interaction images, each video yielded 10,000 images: 9,500 for training and 500 for validation. Because the available candidates depended on the number of retained single-animal blobs and successful donor placements, an excess pool was generated and supplemented as needed before selecting the final 10,000 images. Images generated by clustered pasting were targeted to comprise 5% of the dataset. All samples derived from a given source frame were assigned to the same set, preventing samples from that frame from appearing in both sets. Each selected sample was rotated by 0°, 90°, 180°, or 270° with equal probability.

### Training and detection

Both the label-filtering detector and the detector trained on synthetic interaction images were initialized from pretrained yolo11n-obb weights (https://docs.ultralytics.com/models/yolo11/) and trained on the eight direction classes. Horizontal flip augmentation was disabled, and mosaic augmentation remained active through the final epoch. Both detectors used stochastic gradient descent with an initial learning rate of 0.01 and a final learning rate factor of 1.0. The label-filtering detector was trained for five epochs and the interaction-image detector for 50 epochs. The final-epoch checkpoint was used for each detector. Other Ultralytics augmentation settings remained at their defaults.

For each frame, the detector produced an OBB, confidence score, and direction class for each candidate detection. Detections with confidence scores of at least 0.1 were filtered by the non-maximum suppression implemented in Ultralytics YOLO, applied across direction classes at an OBB IoU threshold of 0.8. When two detections exceeded this overlap threshold, the lower-confidence detection was suppressed. The maximum number of detections per frame was the larger of 300 and twice the known number of animals. Detections were excluded if their assigned class direction was within 22.5° of the OBB short axis. For retained detections, the direction class determined which of the two long-axis directions represented the front. This head direction was recorded as a continuous angle.

### Multi-stage association

Tracking comprised multi-stage association followed by refinement. Association matched detections to a fixed number of track IDs, equal to the configured number of animals (Extended Data Table 2), using previous frames and Kalman filter predictions. Three constant-velocity Kalman filters represented (1) OBB center, (2) OBB shape as width and height, and (3) head direction as a unit vector. Association was initialized at the first frame containing at least the configured number of detections with confidence scores of at least 0.5. The highest-confidence detections, up to the configured number of animals, were assigned sequentially to track IDs. Association proceeded both forward and backward from this frame, and the results were merged into complete trajectories.

Detections were matched to track IDs through six sequential stages (S1–S6) of Hungarian assignment (Extended Data Table 3), using the IoU threshold selected during single-animal blob association. Only unmatched tracks and detections proceeded to the next stage. For S1–S4, the equally weighted cost terms for IoU, angular difference, and consecutive unmatched frames were [(1 − IoU)/(1 − *T*)]^2^, (Δ*θ*/90^∘^)^2^, and *min*(*m*/10,1), respectively, where *T* was the association IoU threshold and *m* was the number of consecutive unmatched frames. For S1 and S3, Δθ was the smaller angle, in degrees, between the reference and detected head directions. For S2 and S4, it was the smaller angle between the reference and detected OBB long axes treating the two opposite orientations of each axis as equivalent. Within each stage, assignment was first performed using the OBB predicted by the Kalman filter and then, for tracks and detections that remained unmatched, using the most recent matched OBB. S5 and S6 used center distance for tracks remaining unmatched after S1–S4. For S5, the reference OBB was the predicted OBB when valid and otherwise the most recent matched OBB. Center distance from this reference was normalized by the square root of the reference OBB area and combined with the same unmatched-frame cost term. The maximum center distance was 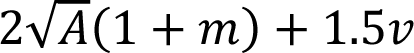, where *A* was the area of the reference OBB and *v* was the Kalman-estimated center speed. S6 applied to tracks unmatched for at least 10 consecutive frames and used the OBB from 10 processed frames earlier as the reference, falling back to the most recent matched OBB when unavailable, with no distance limit.

When S1–S4 or S6 recovered a track after unmatched frames, the intervening frames were rescanned along a linearly interpolated OBB path. In each frame, the nearest detection was accepted if its center was within the larger of twice the square root of the interpolated OBB area and 1.5 times the expected center displacement per frame. S5 recovery did not trigger rescanning, leaving intervening frames unmatched at this stage. Head direction was recorded as missing for S2, S4, S5, and S6 assignments. For S2 and S4, the head-direction Kalman filter was updated using the long-axis orientation closest to the last valid head direction; for S5 and S6, it was not updated. A reversal exceeding 90° and persisting for three consecutive S2 or S4 assignments was retained as a sharp turn. Proximity-based distance assignments were further checked for incorrect recovery. An S5 assignment was removed if the most recent preceding match came from initialization or S1–S4 and the following frame was unmatched. S5 and S6 assignments were also checked retrospectively for transient excursions. If one was followed by another within 10 intervening frames, with no S1–S4 assignment between them, the earlier assignment was removed if the later OBB center was closer than the earlier center to the most recent matched center preceding both assignments. An isolated head-direction reversal in a three-frame window was flipped by 180° if the central direction differed from both neighbors by more than 90° and the flip reduced their summed angular difference.

After merging forward and backward association results, dynamic programming resolved remaining front–rear ambiguities in frames with valid OBBs. Each long axis had two possible directions separated by 180°. The sequence minimizing angular changes relative to available head-direction estimates was selected. If no valid estimate was available before or after a gap in head direction, one direction was selected for the first valid OBB and subsequent directions were chosen to minimize angular changes.

### Refinement

Refinement used geometric and temporal information to fill short gaps and correct positional and head-direction inconsistencies. Internal gaps of fewer than 10 frames were first evaluated for interpolation. Neither bounding OBB could have been assigned by S5 or S6 or generated by earlier interpolation or filling. The OBB after the gap had to meet the association IoU threshold with either the predicted OBB or the OBB before the gap, with a head-direction difference of at most 90° for the qualifying comparison. For eligible gaps, OBBs were linearly interpolated. Head direction was interpolated with angular wraparound accounted for, then assigned to the nearest direction along the interpolated OBB long axis. The entire gap was left unfilled if any interpolated head direction differed from its corresponding long axis by more than 45°.

Remaining gaps of fewer than 10 frames were assessed for overlap with another individual. A gap was attributed to overlap if the same other track overlapped the target immediately before and after the gap, with positive OBB IoU on both sides and IoU of at least 0.1 on at least one side. If multiple tracks qualified, the track with the largest summed IoU before and after the gap was selected. Its OBB was copied into each missing frame for which it was available. The target’s head direction was interpolated with angular wraparound accounted for and assigned to the nearest direction along the copied OBB long axis if the angular difference was at most 45°. Otherwise, the overlapping individual’s head direction was used.

Positional inconsistencies were evaluated when OBBs were available in both adjacent frames. An isolated positional jump was identified when the squared center distance between the preceding and following OBBs was smaller than the sum of squared center displacements into and out of the target frame. The target OBB was replaced by linear interpolation between its neighbors only if this reduced the larger of the two center displacements. Head direction was retained if the target OBB had been assigned at initialization or by S1. Otherwise, head direction was interpolated from adjacent frames with angular wraparound accounted for, then assigned to the nearest direction along the interpolated OBB long axis. The correction was rejected if the angular difference from that axis exceeded 45°.

Short head-direction reversals were corrected by comparison with neighboring sequences. A sequence comprised successive available head directions, each differing from the preceding one by at most 60°. Sequences lasting less than 5 s were examined only if an immediately adjacent sequence lasted at least 5 s and differed in direction by at least 120° at their boundary. All directions in the short sequence were reversed by 180° only if this reduced the summed squared angular difference across the relevant boundaries and connected the sequence to at least one neighboring sequence lasting at least 5 s.

### Optional identity correction using contrastive learning

Identity correction using contrastive learning was adapted from idtracker.ai v6 ^10^. It was enabled when the user indicated that almost complete occlusion was expected and applied at the start of refinement. Training images were obtained from isolated segments: track segments of at least four frames with no OBB overlap with other animals and no change in head direction of 90° or more between consecutive frames. Each frame yielded a grayscale square crop centered on the animal and rotated to point its front upward. Only the connected component of foreground pixels with the greatest overlap with the target OBB was retained; all other pixels were set to black. Crop size was the largest OBB long-side length across isolated segments, rounded up to a multiple of 32 pixels.

Following idtracker.ai v6 ^10^, a ResNet18 network with one input channel and an eight-dimensional output was trained from random initialization. Positive pairs comprised two crops from the same isolated segment; negative pairs came from temporally overlapping segments with different track IDs. The pairwise contrastive loss of Chopra et al. ^25^ penalized distances above 1 for positive pairs and below 10 for negative pairs. The Adam optimizer ^26^ used a learning rate of 0.001. Batches contained equal numbers of positive and negative pairs, with a reference size of 400 pairs of each type, adjusted to available GPU memory. Segment size was the number of frames, with one crop per frame. Sampling weights combined segment-size and loss-score contributions with equal weighting after each was normalized by its sum across eligible segments or segment pairs. The size contribution was the segment size for positive pairs and the sum of both segment sizes for negative pairs. Loss scores were increased for segments or pairs producing nonzero loss, making them more likely to be sampled again. Embedding quality was evaluated by the silhouette score after k-means clustering into as many clusters as the configured number of animals, using up to 1,000 crops per animal. The threshold of 0.91 was adopted from idtracker.ai v6 ^10^. Training stopped after two consecutive evaluations without improvement once the best silhouette score reached 0.91, or after 30 consecutive evaluations without improvement. The checkpoint with the highest score was retained.

An eight-dimensional embedding centroid was calculated for each isolated segment. Identity verification was restricted to interactions with pairwise OBB IoU of at least 0.1, an internal unmatched frame, a filled frame, or the first reappearance after an unmatched sequence. For interactions involving two or three track IDs, the nearest isolated segments before and after the interaction were compared across all participating IDs. Larger interactions were evaluated in subsets of at most three IDs. Each candidate subset comprised a track containing a filled frame or first reappearance, together with the one or two overlapping tracks with the greatest maximum IoU during the interaction. Euclidean distances between embedding centroids formed the Hungarian-assignment cost matrix. Direct verification retained the existing identity assignment only if every matched distance was below 5.0. An alternative assignment was accepted only if its total cost was no greater than half the existing assignment cost and half the next-lowest assignment cost. Otherwise, direct verification was inconclusive.

When direct verification was inconclusive, historical appearance information was used. For each track ID, a prototype was defined as the median of the embedding centroids of all isolated segments ending before the interaction. The pre-interaction segments were first compared with these prototypes under the same acceptance conditions and had to support the existing assignment. Post-interaction segments were then compared with the same prototypes, and identities were corrected only if an alternative assignment met those conditions. For unresolved two-animal interactions, the immediately preceding and following segments were additionally compared with both prototypes. A match required an embedding distance below 5.0 and mutual nearest matching between segments and prototypes. A correction was accepted only if the pre-interaction segment matched its existing ID and the post-interaction segment matched the other ID, unless the other animal independently supported the unchanged assignment.

For each corrected track ID, filled frames were prioritized as relabeling candidates because identity continuity was not supported by direct detections in those frames. The candidate with the greatest OBB IoU with another animal in the interaction was selected. If no filled frame was available, the same criterion was applied to all frames in the interaction. The interaction midpoint was used only if no valid IoU-based candidate existed. The earliest selected frame across corrected IDs became the common relabeling frame.

### Tools compared in the benchmark

AMADEUS was compared with SLEAP (v1.6.1, top-down), DeepLabCut (v3.0.0rc13), idtracker.ai (v6.0.14), UMATracker (Release-15), and UDMT (v1.1.0). For DeepLabCut, a fixed number of frames was annotated per iteration. After training and inference, the same number of outlier frames was extracted, annotated, and added to the training set. Five iterations were performed, except for the 368-ant video, for which one was used. DeepLabCut multi-animal tracking included tracklet stitching, and filtered predictions were evaluated. SLEAP training data were converted from the corresponding DeepLabCut datasets, with a validation fraction of 0.05 and 200 training epochs. SLEAP used the simplemaxtracks tracker with reconnection across single-frame gaps enabled.

For idtracker.ai and UMATracker, foreground segmentation parameters were adjusted by visual inspection for each video to separate animals as clearly as possible. In several challenging videos, segmentation could not consistently isolate each animal’s complete body from other animals and the background.

For UDMT, every animal’s position was manually specified in the first frame. Because the released version did not accommodate the animal counts in some benchmark videos, a patch provided directly by the UDMT authors was applied. The resize coefficient was also adjusted as needed to avoid associated errors. Despite these adjustments, UDMT completed tracking only for the 10-mouse video. In the other five videos, segmentation did not adequately separate animals, and subsequent processing terminated with errors. All other settings and processing steps followed each package’s standard workflow without further modification.

TRex (2.0.0) ^11^ was also tested on all six benchmark videos using both its default pretrained YOLO model and background subtraction, with detection thresholds adjusted for each video. In all six videos, however, many animals were classified as noise or processing terminated with errors, preventing reliable individual tracking. TRex was therefore excluded from the benchmark.

### Benchmark video datasets

To evaluate generality, we used videos differing in species, recording conditions, animal count, body size, interaction frequency, and frequency of forward movement. The videos showed mice, fish, fruit flies, red flour beetles, ants, and interactions between ants and one cricket.

### Mice

We analyzed mice_10_600×686_40Hz_6min.mp4 from the dataset released by Li et al. ^12^ and archived on Zenodo^27^ under Creative Commons Attribution 4.0 International license. The video shows 10 black male mice moving freely in a circular open-field arena, filmed from above under infrared illumination. The arena was 50 cm in diameter and 30 cm high. The original image size of 600 × 686 pixels was retained; the frame rate was 40 fps, and the recording lasted about 6 min (14,331 frames). Mean body length calculated from ground truth (GT) was 64 pixels. The white floor and gray walls produced markedly different foreground contrast as mice attempted to climb the walls. The video also showed frequent overlaps and large shape changes during grooming. For pose estimation training, we labeled the approximate body-axis endpoints (head side as front and tail side as rear) and one auxiliary keypoint on each ear, giving four keypoints per animal. Invisible parts were not labeled. Forty images of 10 animals were labeled in each of five iterations, totaling approximately 8,000 keypoints.

### Fish

We analyzed test.mp4 from the Fish dataset released for the Fish Tracking Challenge 2024 (https://ftc-2024.github.io/, Creative Commons Attribution-NonCommercial-ShareAlike 4.0 International license) ^28^. The video shows 10 sweetfish (*Plecoglossus altivelis*), approximately 7–14 cm long, schooling in a shallow white tank filmed from above. The tank measured 3 m × 3 m, with a water depth of approximately 15 cm ^29^. The original image size of 2,456 × 2,058 pixels was retained; the frame rate was 15 fps, and the recording lasted 11 min 33 s (10,400 frames). Mean body length calculated from GT was 68 pixels. The background included a pale, grid-patterned floor, a central white object, and landmarks along the edges. Surface waves, reflections, intermittent human shadows, and frequent near-complete overlap caused changes in appearance and temporary occlusions. For pose estimation training, we labeled the anterior head tip as front, the tail-side end of the main body, defined by image contrast, as rear, and one auxiliary keypoint near the body center. Invisible parts were not labeled. Forty images of 10 animals were labeled in each of five iterations, totaling approximately 6,000 keypoints.

### Flies

We analyzed a video of 60 female fruit flies (*Drosophila melanogaster*) from the Supplementary Materials of the paper describing YORU, software for behavior classification and assistance with experimental interventions ^30^. The YORU authors provided the video directly with permission for use in this study. Behavior was filmed from above under visible LED illumination in a bowl chamber with sloped walls, 60 mm in diameter and 3.5 mm deep. The region occupied by flies was cropped from the raw AVI file (1,384 × 1,032 pixels, 30 fps) and converted to MP4 at 800 × 800 pixels and 30 fps, reducing file size from 17.9 GB to 203 MB. The recording lasted 5 min (9,000 frames), and mean body length calculated from GT was 35 pixels. Although the low ceiling limited flight, flies frequently overlapped and formed dense aggregates. They appeared dorsal, ventral, or lateral side up and made sudden, rapid movements. For pose estimation training, we labeled the anterior head tip as front, the posterior abdominal tip as rear, and one auxiliary keypoint on each eye. Invisible parts were not labeled. Twenty images of 60 animals were labeled in each of five iterations, totaling approximately 24,000 keypoints.

### Beetles

We analyzed 32 male red flour beetles (*Tribolium castaneum*), ca. 4 mm long, in a shallow circular arena 3D-printed from polylactic acid (PLA) and covered with a transparent acrylic plate. The arena had a flat central floor of radius 25 mm, a shallow outer slope extending to radius 30 mm, and a maximum depth of 1.0 mm. Beetles were refrigerated for 5 min, then placed upright in the arena with forceps. Recording began after all animals resumed walking. The video measured 2,160 × 2,160 pixels at 60 fps and lasted 2 min 47 s (10,000 frames). Mean body length calculated from GT was 151 pixels. Red flour beetles aggregate and frequently move backward, testing label filtering because backward movement violates the assumption that movement direction indicates head direction. For pose estimation training, we labeled the anterior head tip as front, the posterior abdominal tip as rear, and one auxiliary keypoint at each lateral end of the head–thorax boundary. Invisible parts were not labeled. Twenty images of 32 animals were labeled in each of five iterations, totaling approximately 12,800 keypoints.

### Ants

We analyzed 368 ants (*Pristomyrmex punctatus*), ca. 3 mm long, in a circular arena containing an array of cylindrical posts. The shallow arena was 3D-printed from PLA and covered with a transparent acrylic plate. It had a radius of 50 mm, a maximum depth of 0.9 mm, a flat central floor of radius 45 mm, and a shallow outer slope. Posts 0.8 mm in diameter and 0.6 mm high were arranged concentrically at 2.3 mm spacing, allowing ants to walk between them. The video measured 2,160 × 2,160 pixels at 60 fps and lasted 2 min 47 s (10,000 frames). Mean body length calculated from GT was 47 pixels. This benchmark contained the largest number of animals. *Pristomyrmex punctatus* forms dense aggregates, and its head and abdomen are similar in shape; abdominal swelling and bending further alter appearance across frames. For pose estimation training, we labeled the anterior head tip as front and the posterior abdominal tip as rear, giving two keypoints per animal. Invisible parts were not labeled. However, the DeepLabCut model could not detect the ants, and SLEAP required approximately six days of tracking per iteration, so SLEAP was run for only one iteration. Twenty images of 368 animals were labeled, totaling approximately 14,720 keypoints. idtracker.ai, UDMT, and UMATracker terminated with errors and did not complete tracking.

### Ants with one cricket

We analyzed 32 ants (*Tetramorium tsushimae*), ca. 3 mm long, and one ant-loving cricket (*Myrmecophilus* sp.), ca. 3 mm long, which inhabits this ant species’ nests. A circular platform of radius 30 mm was made from a lightly foamed polyvinyl chloride (PVC) plate, with the surrounding area filled with water to the plate’s upper surface. All animals remained on the platform throughout recording. The video measured 2,160 × 2,160 pixels at 60 fps and lasted 2 min 47 s (10,000 frames). Mean body length calculated from GT was 91 pixels for ants and 76 pixels for the cricket. Their similar sizes provided a test of tracking two species simultaneously. For pose estimation, separate datasets and training–inference loops were used for ants and the cricket. For ants, we labeled the anterior head tip as front, the posterior abdominal tip as rear, and auxiliary keypoints at the head–thorax boundary and petiole, giving four keypoints per animal. For the cricket, we labeled the anterior head tip as front, the posterior abdominal tip as rear, and auxiliary keypoints at both lateral ends of the head–thorax boundary, also giving four keypoints. Invisible parts were not labeled. Twenty images of 33 animals were labeled in each of five iterations, totaling approximately 13,200 key points.

### Evaluation metrics

Ground truth (GT) was generated by manually correcting DeepLabCut output for the video of 32 ants with one cricket and SLEAP output for the other videos. Corrections were made in the AMADEUS refinement GUI by repositioning arrows extending from rear to front keypoints. Clear errors were corrected, but small keypoint deviations were not exhaustively corrected. Body length was defined as the mean positive distance between front and rear keypoints across all animals and frames in the GT. An evaluation tolerance of 0.5 body length was used to reduce sensitivity to small GT errors and avoid favoring or penalizing the methods used to initialize GT.

Position and identity tracking were evaluated with py-motmetrics (v1.4.0; https://github.com/cheind/py-motmetrics). For AMADEUS, DeepLabCut, and SLEAP, each animal’s representative position was the midpoint between its front and rear keypoints; for idtracker.ai, UMATracker, and UDMT, the reported coordinates were used directly. Evaluation covered all frames from the first to the last GT frame. In each frame, Euclidean distances were calculated between all GT and estimated positions, excluding pairs more than 0.5 body length apart. The resulting distance matrix was supplied to MOTAccumulator in py-motmetrics. Multiple Object Tracking Accuracy (MOTA) was calculated from the resulting CLEAR MOT events ^22^, and ID F1 score (IDF1) followed the identity metrics framework of Ristani et al. ^24^. Higher Order Tracking Accuracy (HOTA) was calculated at the same single positional threshold of 0.5 body length, rather than averaged across multiple thresholds ^23^. Thus, all three metrics used the same positional matching threshold. The number of identity switches (IDSW) was also calculated using py-motmetrics.

Front and rear keypoint localization was evaluated using Percentage of Correct Keypoints (PCK). For each matched position, distances between estimated and GT front and rear keypoints were evaluated separately. A keypoint was correct if it lay within 0.5 body length of its GT counterpart. PCK was the percentage of evaluated keypoints meeting this criterion. Head direction was evaluated by counting head-direction flips (FLIP). For each matched position, a FLIP was counted when the circular angular difference between the GT and estimated rear-to-front vectors was at least 90°. Unlike PCK, FLIP evaluates which way the body axis points, independently of body size and instantaneous elongation.

### Behavioral analyses

To demonstrate downstream behavioral quantification, we analyzed (i) collective behavior in the 10-fish video, (ii) head direction estimation in the 32-beetle video, and (iii) contact positions during interactions in the video of 32 ants with one cricket. Internal gaps of at most five frames were linearly interpolated. All analyses in this section used AMADEUS results obtained with random seed 0.

### Fish collective behavior

Each animal’s body length was defined as its mean OBB long-side length across valid frames. For each pair in each valid frame, OBB center distance was normalized by the arithmetic mean of the two animals’ body lengths. Pairwise alignment was the inner product of their head-direction unit vectors (cos Δθ, where Δθ was the difference in head direction). The joint density of distance and alignment was estimated on a 128 × 128 grid using a separable Gaussian kernel, with dimension-specific bandwidths determined by Scott’s rule.

Turning events were detected from each animal’s unwrapped head-direction angle. Frames in which the absolute net change over a 0.5 s window reached 45° were grouped into consecutive candidate sequences. Within each sequence, the window with the largest absolute change was selected, and its sign defined the turn direction. Within that window, the event ended at the maximum angular displacement in the turn direction; onset was the last angular minimum in that direction preceding the maximum. Events were retained only if the onset-to-end change reached 45°. Successive event onsets for the same animal had to be at least 0.5 s apart.

For each turning event, the animal turning first was designated the precursor. Another animal was a potential responder if, at precursor turn onset, its center lay within five times its own body length of the precursor’s center. A response was recorded if a potential responder began a turn in the same direction between the next frame and 1.0 s after precursor turn onset. If multiple turns qualified, the earliest was used. For each qualifying pair of turns, we recorded response lag and the precursor’s angular position relative to the responder at precursor turn onset, with 0° indicating the responder’s front. Their joint density was estimated on a 128 × 128 grid using a von Mises kernel for angular position and a Gaussian kernel for response lag. Bandwidths were determined from the data using circular and linear forms of Scott’s rule, respectively.

### Head direction comparisons in beetles

Using the 32-beetle video, we compared the accuracy of head-direction estimates from AMADEUS with direction inferred from centroid tracking. When movement direction is used as a proxy for head direction under the assumption of forward movement, backward movement can result in incorrect direction estimates, and the beetles frequently moved backward. For GT, each animal’s centroid was the midpoint between its front and rear keypoints. GT head direction was defined by the rear-to-front keypoint vector. AMADEUS identities were matched to GT identities at frame 0 by Hungarian assignment using centroid distance, and this fixed correspondence was checked against positional assignment in every frame. Movement direction was inferred from GT centroid displacement over a centered 30-frame interval (0.5 s at 60 fps). Absolute angular errors of AMADEUS head direction and displacement-based direction were calculated relative to GT head direction at the interval midpoint.

Speed was calculated from the same 0.5 s GT centroid displacement, normalized by each animal’s body length at frame 0 and by 0.5 s. To distinguish stopping from moving animals, a Gaussian kernel density estimate was calculated for log_10_ speed. No zero speeds occurred, and non-finite values were excluded before transformation. Density was evaluated at 2,000 equally spaced points between the 0.2nd and 99.8th percentiles. The leftmost local maximum defined the low-speed mode; the highest-density local maximum at a higher speed defined the moving mode. The minimum between these modes yielded a threshold of 0.0216 body lengths per second. Animals were classified frame by frame as stopping below this threshold and moving at or above it. Displacement-based direction was not treated as a head-direction estimate for stopping animals. For moving animals, median angular error was calculated for each individual and compared between AMADEUS and centroid displacement using a paired Wilcoxon signed-rank test.

### Social interactions between ants and one cricket

Each animal’s body length was its mean OBB long-side length across valid frames. For geometric analysis, each OBB was represented by an ellipse with the same center and orientation and with major and minor axis lengths equal to the OBB’s long and short sides, respectively. Contact began in the first frame with overlapping ellipses. It ended when the distance between their boundaries exceeded 0.5 times the ant’s body length for an ant–cricket pair, or 0.5 times the arithmetic mean body length for an ant–ant pair.

The approach interval immediately preceded contact onset and comprised the continuous period during which ellipse-boundary distance remained within one ant body length for ant–cricket pairs or one arithmetic mean body length for ant–ant pairs. Each animal’s net displacement from approach onset to contact onset was projected onto the line directed toward the other animal at approach onset. An approach was unidirectional if only one projection was positive and mutual if both were positive. Cases in which both projections were zero or negative were not observed.

Contact position was determined on the focal ellipse boundary at contact onset. Where the boundaries intersected, arcs of the focal boundary lying inside the other ellipse were identified, and each arc’s midpoint was calculated along the boundary. For a single arc, its midpoint was used. For multiple arcs, the midpoint nearest the other animal’s center in the preceding valid frame was selected. A single boundary intersection was used directly. If no intersection existed, the focal boundary point nearest the other animal’s center in the preceding valid frame was used.

A mean body ellipse was defined for each species from mean OBB dimensions, averaged across all animals and frames for ants and across frames for the cricket. The angular coordinate of each contact-onset position was transferred to the corresponding mean ellipse and expressed as a position along its perimeter. To calculate mean overlap duration, a 360 × 180 grid was defined within the mean ellipse. In each frame with ellipse overlap, grid positions were mapped to the current focal ellipse and tested for inclusion in the other animal’s ellipse. Overlap duration was accumulated at each position and divided by the number of contacts in the map.

Contact-onset density along the ellipse perimeter was estimated using a wrapped Gaussian kernel with bandwidth 0.035 times the mean ellipse perimeter. Contacts were weighted equally for onset density and by their duration in seconds for the corresponding duration distribution. For both estimates, kernel contributions were summed and divided by the number of contacts.

### Software implementation

AMADEUS was implemented in Python and supports versions 3.10–3.12. Dependencies were Ultralytics 8.3.185 (yolo11n-obb), PyTorch 2.7.1, torchvision 0.22.1, OpenCV 4.10.0.84, NumPy 1.26.4, pandas 2.3.3, SciPy 1.11.4, filterpy 1.4.5, scikit-learn 1.7.2, motmetrics 1.4.0, h5py 3.16.0, imageio-ffmpeg 0.6.0, matplotlib 3.10.6, Pillow 11.3.0, psutil 7.1.0, tables 3.10.1, and tqdm 4.67.1. The GUI used customtkinter 5.2.2 and tkinterdnd2 0.6.3; YAML configurations were loaded with PyYAML 6.0.2. Installations with supported NVIDIA GPUs used the corresponding CUDA 12.8 builds. All reported analyses ran on Microsoft Windows.

### Computational hardware

Speed benchmarks used a workstation with an Intel Core i5-12400F CPU, 64 GB RAM, an NVIDIA GeForce RTX 4070 GPU with 12 GB VRAM, a 2 TB WD Blue SN570 SSD, and Microsoft Windows 11 Home. The software also ran on a laptop with an Intel Core i7-1165G7 CPU, 32 GB RAM, and an NVIDIA GeForce GTX 1650 Ti GPU.

### Recommended recording conditions

Because AMADEUS learns directly from input videos, tracking performance depends on recording conditions. It is designed primarily for laboratory videos with a constant number of animals, in which the animals have bodies elongated enough to define an axis and visually distinguishable fronts and rears. We recommend a fixed overhead camera, approximately planar animal motion, stable illumination, an essentially static background, and sufficient contrast, spatial resolution, and frame rate for reliable visual tracking. Animal count should ideally remain constant. Recordings should include sufficient periods with isolated animals to generate training data. Training and analysis videos may be recorded separately under the same conditions.

## Use of generative AI

Generative AI tools assisted software development and manuscript preparation. ChatGPT and Codex (OpenAI) and Claude and Claude Code (Anthropic) assisted implementation of author-specified functionality, computational optimization, and consistency across scripts. Models included GPT-6 and earlier GPT models and Claude Opus 5 and earlier Claude models. The authors reviewed, tested, and validated AI-generated code rather than relying on any single model’s output. ChatGPT and Claude also helped improve the wording and clarity of author-written manuscript text. The authors reviewed and approved all AI-assisted outputs and take responsibility for the final software, analyses, and manuscript.

## Code availability

AMADEUS is available at https://github.com/jpmyrmecol/AMADEUS under the AGPL-3.0-only license.

## Acknowledgements

We thank Hayato M. Yamanouchi, Ryoya Tanaka, and Azusa Kamikouchi for providing the fruit fly videos; Yixin Li for the UDMT patch and, with colleagues, for making the mouse dataset publicly available; Makoto Itoh for confirming that the fish video could be used; and Gonzalo G. de Polavieja for kindly agreeing to the use of aspects of the idtracker.ai approach in AMADEUS. We acknowledge support from the Meiji Institute for Advanced Study of Mathematical Sciences (MIMS), Meiji University, and especially from Hiraku Nishimori.

## Funding

This research was supported by Japan Society for the Promotion of Science Grants-in-Aid for JSPS Fellows (23KJ0665 and 26KJ0327 to Y.N.) and Scientific Research (24H00707 and 26K02091 to S.D.).

## Author contributions

Y.N.: Conceptualization, Methodology, Software, Investigation, Formal analysis, and Writing – original draft. K.M.: Resources, Investigation, and Writing – review & editing. S.D.: Resources, Investigation, Methodology, and Writing – review & editing.

## Competing interests

The authors declare no competing interests.

---

**Extended Data Fig. 1.**
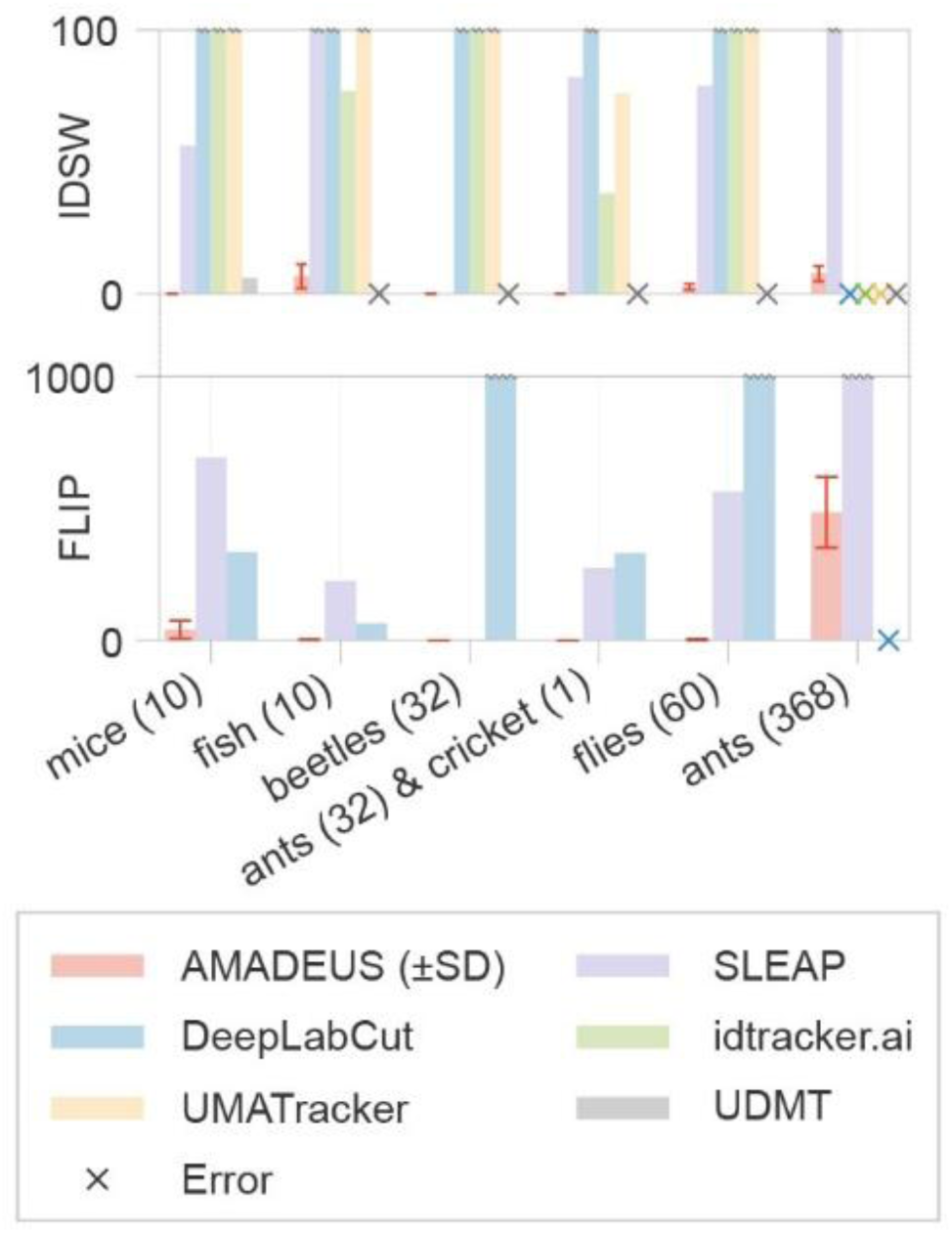
Numbers of identity switches (IDSW) and front–rear flips (FLIP) for six benchmark videos. From left to right, the benchmark videos contain 10 mice, 10 fish, 32 beetles, 32 ants with one cricket, 60 flies, and 368 ants. IDSW and FLIP are shown for the final AMADEUS output after refinement (red), SLEAP (purple), DeepLabCut (blue), idtracker.ai (green), UMATracker (yellow), and UDMT (gray). Post-processing procedures provided by each method were applied where available. Lower values indicate better performance. AMADEUS values show the mean ± standard deviation across three random seeds. Crosses indicate benchmark conditions for which a method failed to produce valid tracking output.

**Extended Data Table 1.** Foreground segmentation settings for benchmark videos. Parameters were adjusted interactively using the real-time preview in the graphical user interface (GUI). In the segmentation mode, bg diff denotes background difference, dark denotes dark-region thresholding, and bg diff + dark denotes background difference intersected with a dark-region mask. Background estimation indicates whether the static background was estimated from the temporal median or maximum at each pixel. Outlier thresholds are the interquartile range (IQR) multipliers defining the lower and upper blob-area bounds as Q1 − k_lower_ × IQR and Q3 + k_upper_ × IQR, respectively. Dashes indicate default settings. For fish, circular ROIs with a radius of 1,228 px were applied to frames 10,288–10,300 and 10,329–10,340, centered at (1,878, 1,029) and (1,228, 1,029), respectively.

| Parameter | Default | Mice<br>(10) | Fish<br>(10) | Beetles<br>(32) | Ants + cricket<br>(32 + 1) | Flies<br>(60) | Ants<br>(368) |
| --- | --- | --- | --- | --- | --- | --- | --- |
| Segmentation mode | bg diff | bg diff + dark | — | — | — | — | dark |
| Background estimation | median | max | — | max | — | max | not used |
| Background difference threshold | 50 | 30 | 70 | 130 | 115 | 70 | not used |
| Intensity threshold | 50 | 45 | not used | not used | not used | not used | 185 |
| Minimum blob area | 10 | — | 20 | — | — | — | — |
| Outlier thresholds | 1.5 / 1.5 | 0.2 / 0.5 | 0.0 / 1.5 | — | 1.0 / 4.5 | 0.0 / 0.5 | — |
| Region of interest (ROI) | not used | — | used | — | — | — | — |

**Extended Data Table 2.** Three questions used to configure downstream processing. Dashes indicate default answers. For the question on overlap severity, Light corresponded to a maximum overlap of 0.2 times the donor mask area, whereas Heavy and Very heavy corresponded to 0.5 times the donor mask area. Very heavy additionally enabled identity correction using contrastive learning during refinement.

| Question | Default | Mice<br>(10) | Fish<br>(10) | Beetles<br>(32) | Ants + cricket<br>(32 + 1) | Flies<br>(60) | Ants<br>(368) |
| --- | --- | --- | --- | --- | --- | --- | --- |
| How many animals are in the video? | 1 | 10 | 10 | 32 | 33 | 60 | 368 |
| How severe is the overlap? | Heavy | — | Very Heavy | Light | — | — | Light |
| Do the animals ever move backward? | Yes | — | No | — | — | — | — |

**Extended Data Table 3.** Stages of multi-stage association. Six sequential stages matched detections to track IDs. Acceptance conditions, assignment costs, and reference OBBs are shown for each stage. Only unmatched tracks and detections proceeded to the next stage. The IoU threshold was selected automatically as described in “Association of single-animal blobs.” In S1–S4, assignment was first performed using the OBB predicted by the Kalman filter and then, for tracks and detections that remained unmatched, using the most recent matched OBB. In S5, the predicted OBB was used as the reference when valid, with the most recent matched OBB used otherwise, and the maximum center distance increased with OBB size, the number of consecutive unmatched frames, and center speed estimated by the Kalman filter. In S6, the OBB from 10 processed frames earlier was used as the reference, with the most recent matched OBB as a fallback when unavailable.

| Stage | Gate | Cost | Reference |
| --- | --- | --- | --- |
| S1 | $\text{IoU} \geq \text{association IoU threshold}$<br>$\text{head direction difference} \leq 90^\circ$ | $\text{IoU difference} + \text{head direction difference} + \text{miss count}$ | KF predicted OBB → last valid OBB |
| S2 | $\text{IoU} \geq \text{association IoU threshold}$ | $\text{IoU difference} + \text{long axis difference} + \text{miss count}$ | KF predicted OBB → last valid OBB |
| S3 | $\text{IoU} \geq 0$<br>$\text{head direction difference} \leq 90^\circ$ | $\text{IoU difference} + \text{head direction difference} + \text{miss count}$ | KF predicted OBB → last valid OBB |
| S4 | $\text{IoU} \geq 0$ | $\text{IoU difference} + \text{long axis difference} + \text{miss count}$ | KF predicted OBB → last valid OBB |
| S5 | adaptive center distance | center distance + miss count | KF predicted OBB (→ last valid OBB) |
| S6 | none | center distance + miss count | OBB from 10 frames earlier |

## Notes

https://github.com/jpmyrmecol/AMADEUS

### Competing Interest Statement

The authors have declared no competing interest.

